# The advent of genome-wide association studies for bacteria

**DOI:** 10.1101/016873

**Authors:** Peter E. Chen, B. Jesse Shapiro

**Author notes:** Corresponding author: B. Jesse Shapiro.

## Abstract

**Highlights:** • The advent of the genome-wide association study (GWAS) approach provides a promising framework for dissecting the genetic basis of bacterial or archaeal phenotypes.

• Bacterial genomes tend to be shaped by stronger positive selection, stronger linkage disequilibrium and stronger population stratification than humans, with implications for GWAS power and resolution.

• An example GWAS in *Mycobacterium tuberculosis* genomes highlights the potentially confounding effects of linkage disequilibrium and population stratification.

• A comparison of the traditional GWAS approach versus a somewhat orthogonal method based upon evolutionary convergence (phyC) shows strengths and weaknesses of both approaches.

**Abstract:** Significant advances in sequencing technologies and genome-wide association studies (GWAS) have revealed substantial insight into the genetic architecture of human phenotypes. In recent years, the application of this approach in bacteria has begun to reveal the genetic basis of bacterial host preference, antibiotic resistance, and virulence. Here, we consider relevant differences between bacterial and human genome dynamics, apply GWAS to a global sample of *Mycobacterium tuberculosis* genomes to highlight the impacts of linkage disequilibrium, population stratification, and natural selection, and finally compare the traditional GWAS against phyC, a contrasting method of mapping genotype to phenotype based upon evolutionary convergence. We discuss strengths and weaknesses of both methods, and make suggestions for factors to be considered in future bacterial GWAS.

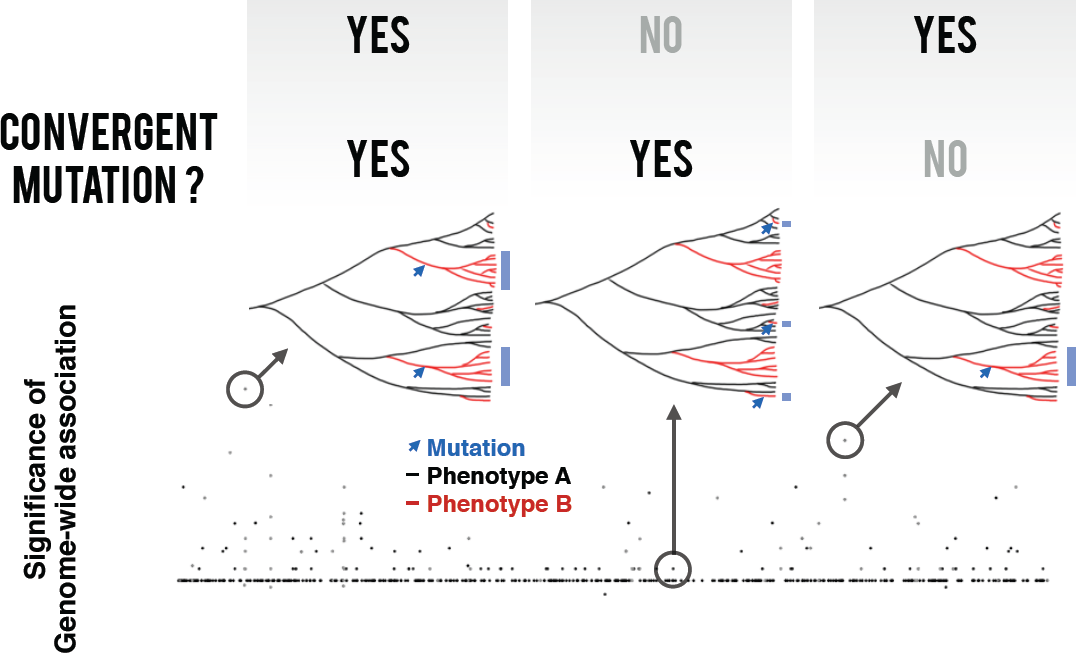

## Introduction

A central goal of biology is to understand how DNA, the primary sequence, gives rise to observable traits. Historically, much effort has gone into deciphering the primary sequence of eukaryotes, primarily *Homo sapiens*. As of August 8, 2014, the National Human Genome Research Institute (NHGRI) reported 1,961 publications of genome-wide association studies (GWAS). Within these studies, a total of 14,014 single nucleotide polymorphisms (SNPs) are associated with over 600 phenotypes. The advent of GWAS in bacteria has mainly occurred in the last two years [1**, 2**, 3**, 4**, 5**, 6**], and provides an unbiased “top-down” framework [7] to dissect the genetic basis of bacterial phenotypes. In principle, any measurable bacterial phenotype (or archaeal phenotype, although here our focus is on bacteria) can be dissected with a GWAS approach. To date, bacterial GWAS have focused on clinically-relevant phenotypes such as virulence and antibiotic resistance, but there is also great potential to investigate environmentally or industrially relevant phenotypes as well.

## Bacterial genomes experience strong linkage, strong stratification and strong selection

Are bacterial genetic mapping studies any different from eukaryotic studies? Although there are many fundamental differences, this review highlights three features that are most germane to GWAS. The impact of the first two differences, in linkage and population stratification, have been recognized before [6**, 7], but we identify the strength of natural selection relative to drift as a third and under-appreciated factor to consider in bacterial GWAS.

First, unlike eukaryotic recombination which occurs predominantly via the crossing-over of two homologous chromosomes during meiosis, bacterial recombination occurs via gene conversion of relatively short stretches of DNA. In bacteria, recombination is not coupled with reproduction, and can occur multiple times within a cell’s lifespan, or not at all. Without any recombination, purely clonal transmission of DNA leaves the entire bacterial chromosome in complete linkage (in strong linkage disequilibrium; LD). As with eukaryotic genomes, bacterial recombination events break this linkage, but the landscape of LD is markedly different from that seen in eukaryotes; gene conversion events leave a “patchwork” of recombined tracts on top of a genomic background of linked regions called a clonal frame [8]. In contrast to eukaryotic LD patterns, all regions of the clonal frame are in complete linkage, and these regions may be quite distant from one another. The clonal frame phenomenon limits the utility of classic genetic mapping methods mainly by obscuring the true causal variant from the rest of the linked sites in the clonal frame. Here, we define a variant as causal if it plays a functional role in the phenotype of interest, as opposed to only being correlated with the phenotype.

Second, as with eukaryotes, bacterial genomic diversity may be shaped by population stratification. Stratification refers to a “situation in which the population of interest includes subgroups of individuals that are on average more related to each other than to other members of the wider population” [9]. These subpopulations give rise to spurious associations when “cases” (with phenotype A) are on average more closely related with each other than with “controls” (without phenotype A); in other words, associations due to genetic relatedness rather than causality for the phenotype of interest. The problem of population stratification is particularly acute in highly clonal (rarely recombining) bacteria, and in those with separate geographic or host-associated subpopulations [6**].

Third, the phenotypes of most interest in bacterial GWAS are largely different from many human disease phenotypes. In particular, bacterial phenotypes tend to be shaped by strong natural selection (e.g. positive directional selection driving drug resistance), while many human disease phenotypes evolve largely by genetic drift owing to historically small effective population sizes (e.g. due to population bottlenecks); in this scenario, drift overpowers purifying selection and leaves slightly deleterious alleles in the population that underlie disease traits [10, 11]. This is not to say that bacteria do not experience genetic drift (particularly in frequently bottlenecked populations), but simply that many traits of interest (e.g. resistance, virulence, host-association) have evolved recently and under strong positive selection. These bacterial traits might also be controlled by mutations with large effect sizes on the phenotypes of interest. If this is the case, relatively small samples of bacterial genomes should be sufficient to identify causal mutations [11, 12].

## Units of genetic and phenotypic variation

The two basic requirements for GWAS are genotypic and phenotypic measurements from a sample of organisms. Phenotypes are usually broken into either discrete units (e.g. resistance/sensitive or high/low virulence) or continuous traits (e.g. human height). Phenotypes must be reproducible, and easy to measure, ideally in high-throughput if hundreds or thousands of samples are being studied. At the genotypic level, a set of bacterial genomes can be broken down into a “core” genome shared among nearly all members and an “accessory” genome composed of elements present in some strains but not others (typically including genes involved in environmental adaptation) [13, 14]. The genetic units of a GWAS may be variants in the core (e.g. single nucleotide polymorphisms (SNPs) or small indels) [2**, 3**, 4**, 5**] or in the flexible genome (e.g. presence/absence of larger pieces of DNA including genes or operons [1**, 15, 16, 17] (Table 1). While most bacterial GWAS to date have studied either SNPs or gene presence/absence, Sheppard et al. [1**] described a method that uses n-mers (“words” of DNA) as the basic unit of association, allowing them to study both the core and flexible genome simultaneously.

**Table 1.**

| Study | Year | Taxa | Relative recombination rate | # genomes | Phenotype | Association method | Addresses accessory genome? | Unit of genetic variation studied | # of variants | Correction for population stratification |
| --- | --- | --- | --- | --- | --- | --- | --- | --- | --- | --- |
| Sheppard et al. [1] | 2013 | *C. jejuni* | moderate | 29 (+ validation in 161) | host specificity | allele counting | yes | 30-bp DNA sequences (words) | >10,000 words (?) | simulation of word gain/loss along the phylogenetic tree |
| Farhat et al. [2] | 2013 | *M. tuberculosis* | low | 123 | antibiotic resistance | homoplasy counting | no | SNPs | ~25,000 | implicit in phylogenetic convergence criterion |
| Chen & Shapiro (This review) | 2015 | *M. tuberculosis* | low | 123 | antibiotic resistance | allele counting | no | SNPs | ~3,000 MAF > 0.05) | inferred ancestry clusters |
| Laabei et al. [3] | 2014 | *S. aureus* | low | 90 | virulence | allele counting | no | SNPs & small indels | ~3000 | genomic control |
| Alam et al. [4] | 2014 | *S. aureus* | low | 75 | antibiotic resistance | allele counting and homoplasy counting | no | SNPs | ~55,000 | inferred ancestry clusters |
| Chewapreecha et al. [5] | 2014 | *S. pneumoniae* | high | 3085 (+ validation in 616) | antibiotic resistance | allele counting | no | SNPs | ~400,000 (MAF > 0.01) | inferred ancestry clusters |
| Salipante et al. [16] | 2014 | *E. coli* | low-moderate | 312 | antibiotic resistance | allele counting | yes | gene presence/absence | ~15,000 genes | inferred ancestry clusters |
| Chaston et al. [17] | 2014 | 41 strains | N/A | 41 | host development time and triglyceride content | allele counting | yes | gene presence/absence | ~12,000 genes | consideration of genes with unique phylogenetic distributions |
| van Hemert et al. [15] | 2010 | *L. plantarum* | low | 42 | host immune response | allele counting | yes | gene presence/absence | ? (CGH) | none |

SNP = single nucleotide polymorphism, MAF = minor allele frequency, CGH = comparative genomic hybridization

## Allele counting and homoplasy counting approaches to GWAS

GWAS approaches for bacteria can be broadly broken down into allele counting [1**, 3**, 4**, 5**] and homoplasy counting [2**, 12] methods (Table 1 and Graphical Abstract). The primary association signal for allele counting methods is derived from an over-representation of an allele at the same site in cases relative to controls, which can later be corrected for population stratification. In contrast, homoplasy counting methods (in this case, phyC [2**]) derives its evidence of association by counting repeated and independently emerged mutations occurring more often on branches of cases relative to controls. Homoplasy, as an indicator of convergent evolution, is a well-known signal of positive selection [28]. Combining this signal of selection with phenotypic associations (e.g. convergent mutations that occur only in cases and not in controls) provides the basis for homoplasy-based association tests.

## Architecture of a strong association signal

GWAS signals from allele counting and homoplasy counting methods are not expected to perfectly overlap because each method represents different strengths and weaknesses. However, with a sufficiently large sample size, allele counting methods theoretically can detect all convergent sites (identified by homoplasy counting methods) as well as non-convergent sites. Still, ever-increasing sample size does not directly address the confounding effects of both population stratification and LD on allele counting methods. Homoplasy counting intrinsically accounts for these effects by virtue of its phylogenetic convergence criterion. In contrast, allele counting methods have no such phylogenetic requirement. Thus, a monophyletic group containing many cases with the same over-represented allele at the same site may provide a strong signal for allele counting while providing no signal for homoplasy counting. Conversely, homoplasy counting requires a smaller count of homoplasy events (versus allele counts) in order to reach statistical significance; thus, a relatively small sample size with a strong paraphyletic structure may provide homoplasy counting with a much stronger signal than allele counting.

## A genome-wide association study of antibiotic drug resistance in *Mycobacterium tuberculosis*

To examine the impacts of clonal frames (strong LD) and population stratification, we performed a ‘traditional’ GWAS using PLINK on a population of 123 *M. tuberculosis* (MTB) genomes that had been previously analyzed by phylogenetic convergence (phyC) [2**]. Of the 123 strains, 47 (cases) are resistant to at least one antibiotic and 76 strains are sensitive to all antibiotics (controls). This dataset contains 11 ‘gold standard’ experimentally-verified antibiotic resistance alleles, all of which were identified by phyC, along with 39 new phyC hits in nonsynonymous coding sites and intergenic regions, and 7 hits in synonymous sites. We chose this particular MTB dataset as it allows a comparison of the results from traditional GWAS and phyC, and also because MTB genomes possess extensive LD and strong population structure, making them challenging subjects for traditional GWAS.

## Clonal frames and the resolution of GWAS signals

MTB is considered to be a highly clonal pathogen, with very little detectable recombination [18]. Consistent with this, we observe a clonal frame consisting of linked sites across the genome. This clonal frame is evident from the extensive genome-wide linkage (black or red in Figures 1a and b, respectively), interrupted by a few homoplasic sites (small white or black points, respectively) identified by the four-gamete test [19] or the D’ measure [20] of linkage (Figure 1a and b). The r^2^ measure [21] does not directly measure recombination or homoplasy, but rather how well the allelic state at one site in the genome can predict the allele present at another site. The r^2^ analysis confirms that MTB has extensive genome-wide LD, posing a challenge to pinpointing causal variants (Figure 1c and d). Other more highly recombining bacteria, such as *Streptococcus pneumoniae* (Figure 1e) have less long-range LD and more localized, shorter LD blocks (black triangles near the horizontal axis), facilitating GWAS [5**]. Because the extent of genome-wide linkage is unlikely to be known *a priori*, an important first step before performing a bacterial GWAS is to characterize LD, as illustrated here (Figure 1).

**Figure 1.**
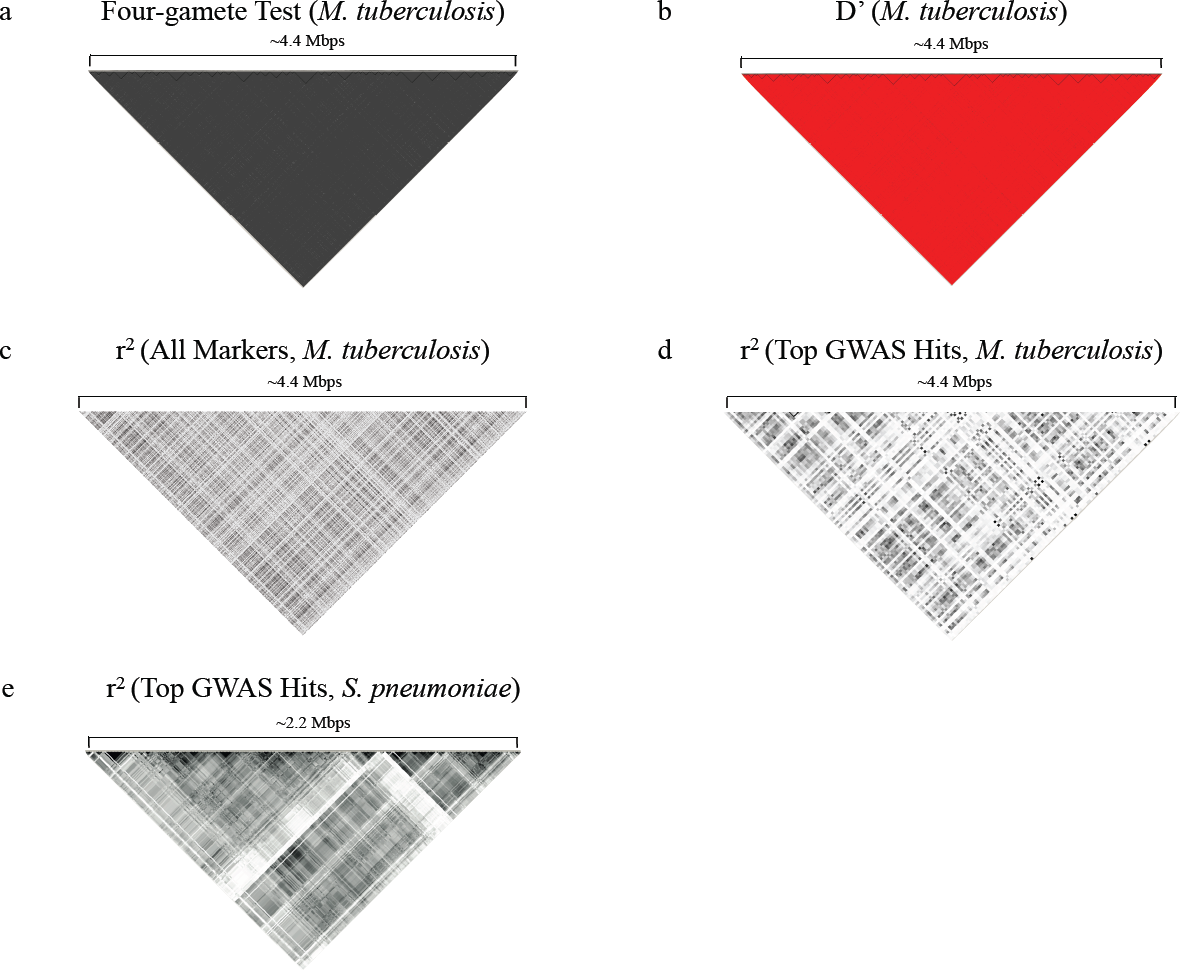
Patterns of linkage disequilibrium (LD) in bacterial genomes assessed by different metrics. The x-axis of heat maps (a-d) represents the physical position along the MTB genome; (e) shows the *S. pneumoniae* genome. Each square in the heat map represents a pairwise calculation of LD. a) Four-gamete test. White squares denote four observed haplotypes indicating recombination may have occurred between the two sites. Black squares denote three or fewer observed haplotypes (strong linkage). b) Pairwise |D’| measurements (range of |D’| values: 0 ≤ |D’| ≤ 1). Red squares denote |D’| = 1 (strong linkage). Black squares denote |D’| < 1. c) Pairwise r^2^ measurements (range of r^2^ values: 0 ≤ r^2^ ≤ 1). Black squares denote r^2^ = 1 (strong correlation). The lighter squares denote progressively smaller r^2^ values. d) Pairwise r^2^ measurements for the top 133 GWAS hits only. Black squares denote r^2^ = 1. The lighter squares denote progressively smaller r^2^ values. e) Pairwise r^2^ measurements of beta-lactam resistance associated variants co-detected in two separate *S. pneumoniae* populations [5**]. Black squares denote r^2^ = 1. The lighter squares denote progressively smaller r^2^ values.

## Correcting for population stratification

The strong clonal nature of MTB also creates strong population substructures that in turn may lead to false positive associations. Without any population stratification correction we observe a substantial systematic inflation of the association test p-values (Figure 2a), likely due to both causal and non-causal resistance-associated mutations being linked on the same clonal frame. We assessed two classic methods of addressing population stratification. The first method, called genomic control [22], normalizes all p-values by a single inflation factor λ, which is the observed median chi-square divided by the expected median chi-square with 1 degree of freedom. Due to a relatively large observed inflation factor (λ = 12.20), genomic control seems to over-correct, leaving no statistically significant GWAS hits (Figure 2b). A less conservative correction for population stratification is to infer ancestry by identifying genetic subpopulations within the overall population, and then testing for association conditional on these subpopulations. Subpopulations can be inferred using a variety of methods (e.g. multi-dimensional scaling in PLINK [23], principal component analysis in EIGENSTRAT[24], and BAPS [25]), then used as covariates in association testing (e.g. with the Cochran-Mantel-Haenszel test). Here, we defined subpopulations based on 14 previously defined MTB epidemiological clusters [2**]. Using these epiclusters as covariates reduced the inflation factor to 1, suggesting that it effectively controls for population stratification (Figure 2c). Although this procedure clearly changes the Manhattan plot (Figure 2, right panels), producing more clean ‘hits’ that stand out from the average p-value, we note that none of these hits pass correction for multiple hypothesis testing. Therefore, correcting for population stratification can reduce GWAS power significantly – a problem that could potentially be overcome by using larger sample sizes (e.g. thousands rather than hundreds of genomes; [5**]).

**Figure 2.**
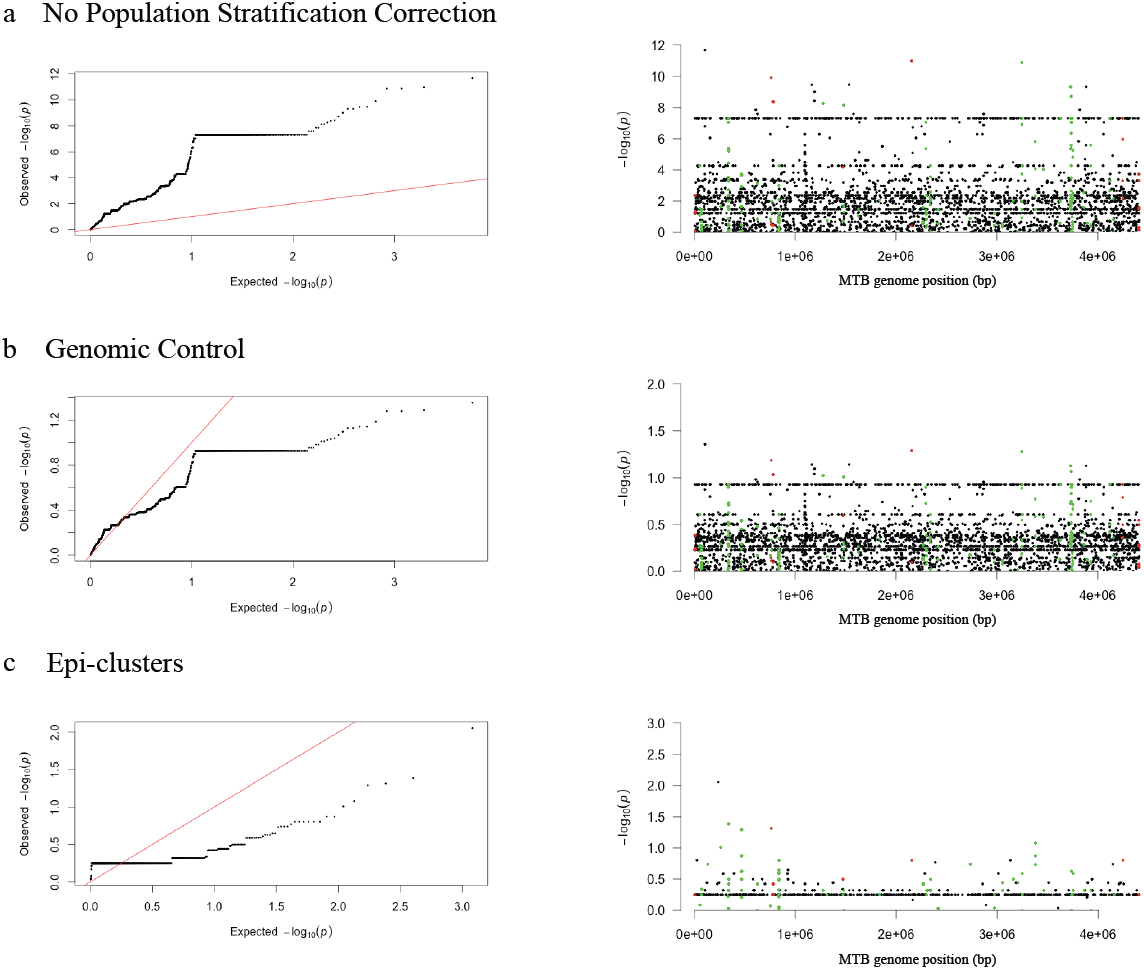
GWAS for antibiotic resistance in MTB. GWAS was performed using Plink version 1.07 [23]. The x=y line (red in QQ plots; left) represents the null hypothesis of no association signal. In Manhattan plots (right), SNPs in ‘Gold Standard’ resistance genes are shown in red, and SNPs in phyC candidate genes in green (excluding synonymous sites). Different corrections for population stratification were applied: a) No population stratification correction. b) Population correction with genomic control. c) Population correction using “epi-clusters” and Cochran-Mantel-Haenszel 2x2xK test, where K = 14 epi-clusters.

## Comparison of GWAS against convergence testing

Despite the lack of significance after multiple testing correction, we identified 133 potential GWAS hits (SNPs) in 77 genes that stood out as outliers from the average genome-wide p-value (Figure 2c), which we will discuss for illustration purposes. These GWAS hits overlapped with 5 of 11 previously known ‘gold standard’ resistance genes and 22 of 46 additional phyC candidate resistance genes. It is also evident that correcting for population stratification improves the overlap with known resistance genes and phyC hits (Figure 3).

**Figure 3.**
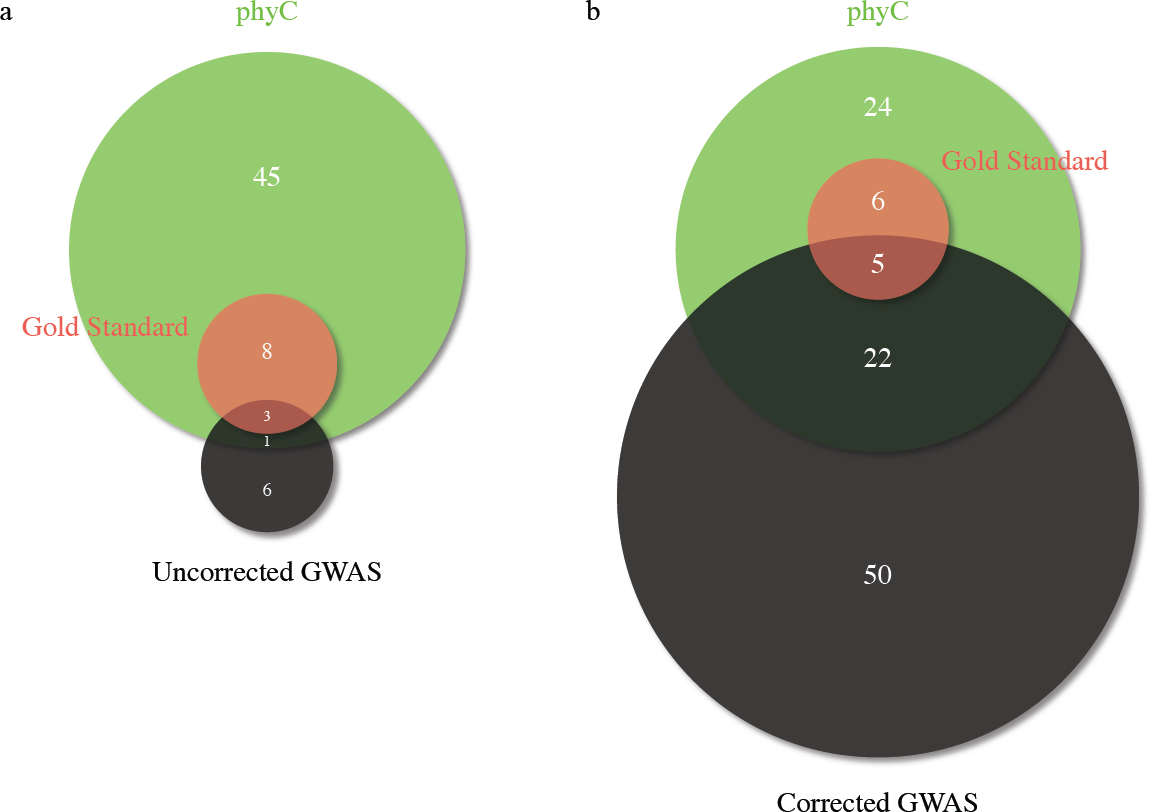
Overlap among GWAS candidates, phyC candidates, and “Gold standard” resistance genes. Numbers in Venn diagrams are in units of genes or intergenic regions. a) No population stratification correction. b) Population correction using “epi-clusters” and Cochran-Mantel-Haenszel 2x2xK test, where K = 14 epi-clusters.

## Potential new drivers of drug resistance

Of these top 133 GWAS hits, 75 SNPs (in 50 genes) did not overlap with either known resistance genes nor with phyC candidates (Figure 3b). Due to long-range LD, it is not immediately clear without further analysis whether these 75 SNPs represent false positives due to their correlation with the true drivers of resistance, either ‘gold standard’ resistance genes or phyC candidates. However, 15 out of these 75 SNPs were relatively uncorrelated (r^2^ < 0.3) with any of the other 133 top GWAS hits, suggesting they could play causal roles in resistance phenotypes. As an example to illustrate the importance of assessing LD patterns around GWAS hits, the top GWAS hit (a nonsense mutation in an oxidoreductase gene, Rv0197) can be viewed from two different perspectives:

1. The top GWAS hit may be a false positive because it is in moderate correlation (0.4 < r^2^ < 0.5) with two phyC candidates (PPE9 and PE_PGRS4 genes) and two other GWAS hits (PE-PGRS30 and PE-PGRS46 genes), and does not represent a true causal variant.
2. The top GWAS hit may be driving the association. Although it is in moderate correlation with four other phyC or GWAS hits, all four hits reside within the PE/PGRS families of genes, which are highly polymorphic and might represent false positive associations [2**].

Whether this GWAS hit is causal or not can only be firmly established with followup experiments.

## Future Directions

We have shown the potential of GWAS for bacterial genomes while highlighting two key obstacles: long-range LD within the clonal frame and extensive bacterial population stratification both reduce our ability to pinpoint causal mutations with confidence. However, a third feature of bacterial genomes – the relative strength of positive selection – provides an opportunity to increase the resolution and confidence of GWAS hits. One could combine positive selection tests and GWAS, as has been done previously for traits shaped by positive selection [26, 27]. This approach may potentially address the problem of clonal frames obscuring true causal variants and making them indistinguishable from linked non-causal variants. This idea attempts to identify causal variants in two steps:

1. perform a genome-wide selection scan identifying any genomic regions that are putatively under positive selection
2. perform a “targeted” association study on each genomic region under positive selection

The rationale here is that each genomic region identified as being under positive selection effectively “unlinks” the putative causal variants from its background clonal frame, provided that the selection test itself can distinguish a positive selection region from the clonal frame upon which it occurred [28]. Since positive selection alone does not provide sufficient evidence that a region is associated with the phenotype of interest, step two targets each of the genomic regions identified in the selection scan and tests each one for association with a phenotype of interest. In phyC, the two steps are done simultaneously, using convergence as the signal of positive selection and the *specificity* of convergence to cases but not controls as the association signal. Future work might ‘mix and match’ different signals of selection and association.

As this new and growing field develops, we envision a future where multiple genetic mapping approaches – including GWAS, phyC and selection scans – are combined. Each method may harbor its own strengths and weaknesses so that when combined, each method provides distinct information, thus increasing the power to detect true and causal associations.

## Acknowledgements

This work was funded by the Natural Sciences and Engineering Research Council, the Canadian Institutes for Health Research and the Canada Research Chairs program. We would like to thank Luis Barreiro and Jean-Baptiste Leducq for valuable feedback and discussions.

